# Dissecting causal pathways using Mendelian randomization with summarized genetic data: application to age at menarche and risk of breast cancer

**DOI:** 10.1101/160663

**Authors:** Stephen Burgess, Deborah J Thompson, Jessica MB Rees, Felix R Day, John R Perry, Ken K Ong

## Abstract

Mendelian randomization is the use of genetic variants as instrumental variables to estimate causal effects of risk factors on outcomes. The total causal effect of a risk factor is the change in the outcome resulting from intervening on the risk factor. This total causal effect may potentially encompass multiple mediating mechanisms. For a proposed mediator, the direct effect of the risk factor is the change in the outcome resulting from a change in the risk factor keeping the mediator constant. A difference between the total effect and the direct effect indicates that the causal pathway from the risk factor to the outcome acts at least in part via the mediator (an indirect effect). Here, we show that Mendelian randomization estimates of total and direct effects can be obtained using summarized data on genetic associations with the risk factor, mediator, and outcome, potentially from different data sources. We perform simulations to test the validity of this approach when there is unmeasured confounding and/or bidirectional effects between the risk factor and mediator. We illustrate this method using the relationship between age at menarche and risk of breast cancer, with body mass index (BMI) as a potential mediator. We show an inverse direct causal effect of age at menarche on risk of breast cancer (independent of BMI) and a positive indirect effect via BMI. In conclusion, multivariable Mendelian randomization using summarized genetic data provides a rapid and accessible analytic strategy that can be undertaken using publicly-available data to better understand causal mechanisms.

## Introduction

Mendelian randomization is the use of genetic variants as instrumental variables to assess and estimate the causal effect of a risk factor on an outcome [Davey Smith and Ebrahim, 2003; Burgess and Thompson, 2015b]. A risk factor has a causal effect on an outcome if intervening on the risk factor leads to changes in the outcome. Correlation between a risk factor and an outcome may arise because the risk factor is a cause of the outcome. However, it may also reflect confounding (the risk factor and outcome have common causes) or reverse causation (the outcome is a cause of the risk factor). Instrumental variable analysis represents one way of assessing whether there is a causal effect of the risk factor on the outcome under certain assumptions using observational data.

For a genetic variant to be a valid instrumental variable, it must satisfy three assumptions. First, the genetic variant must be associated with the risk factor. Secondly, the genetic variant must not be associated with confounders of the risk factor-to-outcome association. Thirdly, the genetic variant must not affect the outcome except via the risk factor of interest (no direct effect on the outcome) [Greenland, 2000; Lawlor et al., 2008]. Whereas phenotypic variables tend to display widespread correlations with other phenotypes, genetic variants are often more specific in their associations [Davey Smith et al., 2007], meaning that Mendelian randomization investigations are less susceptible to biases from confounding that adversely affect observational studies. Additionally, as the genetic code is fixed at conception, genetic associations are less susceptible to reverse causation or confounding due to environmental factors.

The instrumental variable assumptions can be assessed to some extent by testing for as-sociations between the genetic variants and potential measured confounders [Burgess et al., 2015b]. However, it is possible that a covariate associated with a genetic variant is not a confounder, but rather a mediator on the causal pathway from the risk factor to the outcome [Haycock et al., 2016]. This is particularly likely if several variants all have directionally con-cordant associations with the same covariate. Genetic associations with a mediator may not represent pleiotropic effects of the variants, but rather represent downstream consequences of intervening on the risk factor. In such a case, the genetic variants are still valid instru-ments, as the only causal pathway from the variants to the outcome is via the risk factor (and potentially also via the mediator).

In many scenarios, it is relevant not only whether the risk factor is a cause of the outcome, but also via what mechanism this causal effect acts. Mediation analysis can be used to dissect the total causal effect of the risk factor on the outcome into an indirect effect of the risk factor on the outcome via the mediator, and a direct effect of the risk factor on the outcome not via the mediator (possibly via other causal pathways or other mediators) [VanderWeele and Vansteelandt, 2009]. This is illustrated in Figure 1. The total effect is defined as the change in the outcome resulting from intervening on the risk factor (say, increasing its value by 1 unit). The direct effect is the change in the outcome resulting from intervening on the risk factor but holding the mediator constant. The indirect effect is the change in the outcome resulting from manipulating the value of the mediator as if we had intervened on the risk factor, but in fact holding the risk factor constant. If all variables are continuous and all relationships between variables are linear, then the total effect is equal to the direct effect plus the indirect effect. Formally, a direct effect defined by intervening on the risk factor and mediator separately is a controlled direct effect, which does not have a counterpart indirect effect. If all relationships are linear, then the controlled direct effect is equal to the natural direct effect, which does have a counterpart, the natural indirect effect. Full details are provided in the Supplementary Material A.1.

**Figure 1:**
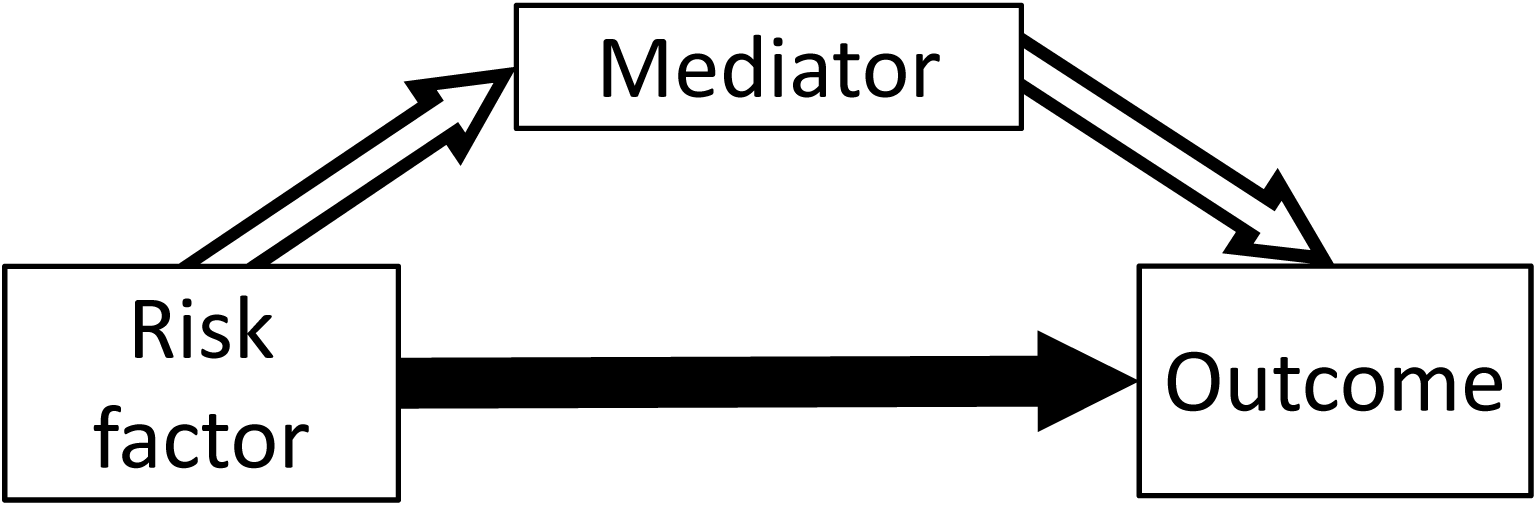
Total effect of risk factor on outcome comprises an indirect effect (hollow arrows) via mediator, and a direct effect (solid arrow) via other pathways.

[Figure 1 should appear about here.]

Mendelian randomization analyses using summarized data have recently become widespread due to the increasing public availability of suitable data in large sample sizes from GWAS consortia, and the possibility of ‘two-sample’ Mendelian randomization in which genetic as-sociations with the risk factor and outcome are estimated in different samples [Burgess et al., 2015b]. It has previously been demonstrated that a (univariable) Mendelian randomization estimate can be obtained from summarized data (beta-coefficients and standard errors) by regressing genetic associations with the outcome on genetic associations with the risk factor [Burgess et al., 2016]. This represents the total effect of the risk factor on the outcome. It has also been demonstrated that direct causal effects of related risk factors can be estimated by regressing genetic associations with the outcome on genetic associations with each of the risk factors in a multivariable regression model; this is referred to as multivariable Mendelian randomization [Burgess and Thompson, 2015a].

In this report, we demonstrate how the total effect and the direct effect of the risk factor on the outcome can be estimated from summarized data, we consider the assumptions necessary for genetic variants to satisfy for consistent estimation, and we exemplify how these estimates can be used to interrogate causal mechanisms with an applied example of the effect of age at menarche on breast cancer risk, with body mass index (BMI) as a potential mediator.

## Methods

### Assumed framework of summarized data and genetic associations

We initially assume that all variables are continuous, and relationships between variables (in particular, the genetic associations with the risk factor *X*, mediator *M*, and outcome *Y*, and the causal effects of the risk factor and mediator on the outcome, and of the risk factor on the mediator) are linear with no effect modification (that is, they are the same for all individuals in the population and do not vary for different values of the independent variable). For each genetic variant *G*_*j*_ (*j* = 1, 2*,…, J*), we assume that we have an estimate 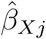 of the association of the genetic variant with the risk factor obtained from linear regression. Similar association estimates are assumed to be available for the mediator 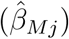 and outcome 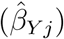. The standard error of the association estimate with the outcome is 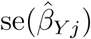. If any of the variables is binary, then these summarized association estimates may be replaced with association estimates from logistic regression; more detail on the binary outcome case is provided later in the paper. The relationships between these variables are illustrated in Figure 2.

**Figure 2:**
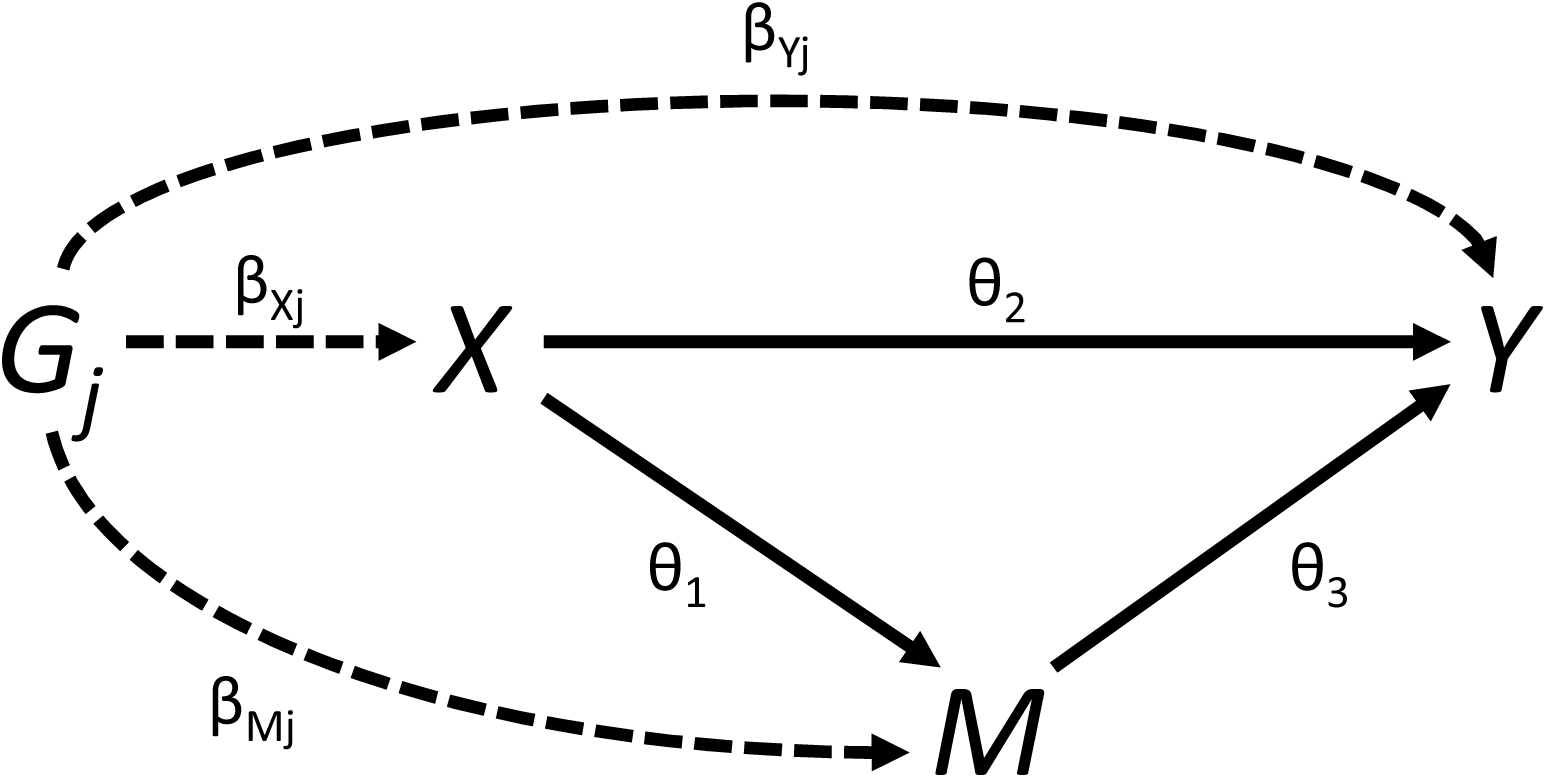
Graphical diagram of relationships between risk factor (*X*), mediator (*M*), outcome (*Y*) and genetic variant (*G*_*j*_). Causal relationships between variables are indicated by solid lines. Associations of the genetic variant are indicated by dashed lines. The direct effect *θ*_*D*_ = *θ*_2_. The indirect effect *θ*_*I*_ = *θ*_1_*θ*_3_. The total effect *θ*_*T*_ = *θ*_*D*_ + *θ*_*I*_ = *θ*_2_ + *θ*_1_*θ*_3_.

[Figure 2 should appear about here.]

We also assume that all genetic variants are uncorrelated (that is, not in linkage disequilibrium). Although conventional instrumental variable methods for analysing summarized data from correlated variants have been developed [Burgess et al., 2016] and software code for analysing correlated variants is provided in the Supplementary Material, as we shall see later there are problems of identification in the mediation setting that may be accentuated by the use of correlated variants. Although this is a strict assumption, often genetic variants in Mendelian randomization investigations are chosen to be the top hits from different gene regions identified by a genome-wide association study, and so the assumption is naturally satisfied. The method makes no specific requirements for the level of statistical significance of the associations between the genetic variants and the risk factor, but variants with robustly verified associations represent more informative instrumental variables.

### Weighted regression for estimation of total and direct effects

If 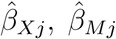, and 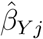 are the genetic associations of variant *G*_*j*_ (*j* = 1, 2*,…, J*) with the risk factor (*X*), mediator (*M*) and outcome (*Y*), and se (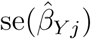) are the standard errors of the genetic associations with the outcome, then the weighted regression:

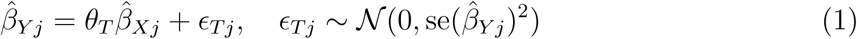

provides an estimate of the total effect of the risk factor on the outcome *θ*_*T*_, known as the inverse-variance weighted estimate [Burgess et al., 2013]. This regression model does not take into account uncertainty in the genetic associations with the risk factor; however, these associations are typically more precisely estimated than those with the outcome, and ignoring this uncertainty does not lead to inflated Type 1 error rates in realistic scenarios [Burgess et al., 2013].

The inverse-variance weighted estimate can be motivated as the fixed-effect meta-analysis pooled estimate of the variant-specific causal estimates 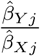 with standard errors taken as 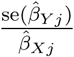 (the leading order term from the delta expansion for the standard error of the ratio of two variables). This meta-analysis estimate can also be obtained by the weighted regression model in equation 1 [Thompson and Sharp, 1999]. The weighted regression model can be expanded by including genetic associations with the mediator:

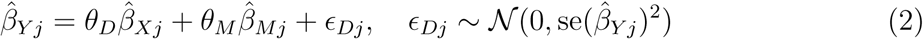

to provide an estimate of the direct effect *θ*_*D*_. The weighted regression method for calculating the total effect (equation 1) is equivalent to the two-stage least squares (2SLS) method with individual-level data, in which the first stage of the method regresses the risk factor on the genetic variants, and the second stage regresses the outcome on fitted values of risk factor [Burgess et al., 2016]. The weighted regression method for calculating the direct effect (equation 2) is also equivalent to a two-stage regression method, except that the first stage also regresses the mediator on the genetic variants, and the second stage regresses the outcome on fitted values of the risk factor and fitted values of the mediator [Burgess et al., 2015a]. Software code to implement these analyses is provided in Supplementary Material A.2. With a continuous outcome, the indirect effect of the risk factor on the outcome can be calculated as *θ*_*I*_ = *θ*_*T*_ *-θ*_*D*_.

For consistent estimation, it is required that all genetic variants used to estimate the total effect of the risk factor on the outcome satisfy the standard assumptions of Mendelian randomization: they are associated with the risk factor, not associated with confounders, and there is no pathway from any genetic variant to the outcome except via the risk factor. All variants used to estimate the direct effect of the risk factor on the outcome must satisfy the assumptions of multivariable Mendelian randomization: they are associated with the risk factor and/or mediator, not associated with confounders, and there is no pathway from any genetic variant to the outcome except via the risk factor and/or the mediator [Burgess and Thompson, 2015a].

### Identification of the direct effect

If the genetic associations with the mediator are entirely determined by their associations with the risk factor, then with an infinite sample size (if associations are perfectly linear with no heterogeneity) the direct effect would not be identified, as the genetic associations with the risk factor and mediator would be perfectly correlated. Hence, it is necessary for there to be some heterogeneity in the genetic associations or the relationships between the variables. This may occur for a complex variable such as BMI, where different genetic variants may influence BMI in different ways or via different biological pathways, potentially leading to different magnitudes of causal effect on the mediator and/or outcome. Alternatively, if there are genetic variants that are instrumental variables for the mediator only, then these variants could be included in the multivariable Mendelian randomization analysis. However, such variants are not valid instrumental variables for the risk factor, and so should not be used to estimate the total causal effect of the risk factor on the outcome.

### Applied example

As an illustrative example, we consider the causal effect of age at menarche on breast cancer risk. Numerous genetic variants have been discovered that influence age at menarche. Later puberty reduces the total number of ovulatory cycles and hence the life-time sex-hormone exposure, thus we expect later menarche to be protective for breast cancer. This is in line with observational epidemiological findings [Collaborative Group on Hormonal Factors in Breast Cancer, 2012]. However, later menarche is also associated with lower BMI, and it is known that genetically predicted BMI (and also adolescent BMI) is inversely associated with breast cancer risk [Guo et al., 2016; Baer et al., 2010]. Therefore age at menarche will likely have an indirect effect on breast cancer risk via BMI as well as a direct effect (in the opposite direction) not via BMI.

We have taken 375 genetic variants demonstrated to be associated with age at menarche at a genome-wide level of significance [Day et al., 2017]. Genetic associations with age at menarche (measured in years) were obtained from the Reprogen consortium based on 329,000 women of European descent. Genetic associations with BMI were obtained from the GIANT consortium, based on 339,000 individuals, 95% of whom are of European descent [Locke et al., 2015]. Genetic associations with breast cancer risk were obtained from the Breast Cancer Association Consortium (BCAC) on 47,000 cases and 43,000 controls (all female) of European descent [Michailidou et al., 2015]. Although genetic associations with BMI were estimated at different timepoints for different studies in the GIANT consortium, as genetic variants typically influence variables across the whole life-course, it is not crucial when these associations are measured, provided that they are measured in individuals before they have disease events (to prevent reverse causation, see Discussion for more detail). A more detailed analysis of these same data (although based on the individual-level data) was previously reported by Day et al. [2017]; further details relating to applied aspects of the analysis are provided in that paper.

Univariable Mendelian randomization suggested a null effect of age at menarche on breast cancer risk (odds ratio per 1 year later menarche 1.00, 95% confidence interval 0.96, 1.05). However, a multivariable Mendelian randomization analysis adjusting for genetic associations with BMI suggested a protective direct effect of later age at menarche (odds ratio 0.94, 95% confidence interval 0.89, 0.98). This suggests that an intervention to delay menarche would have no net effect on breast cancer risk if it also had the expected consequence of lowering adolescent BMI (or, similarly, if the delay in menarche was achieved by reducing pre-pubertal BMI). However, an intervention which had an effect on post-pubertal sex-hormone exposure equivalent to a later menarche would be likely to have a protective effect on breast cancer risk, as such an intervention could not affect pubertal timing and hence would not alter BMI; hence only the direct effect of age at menarche on breast cancer risk would apply here. We note that the results presented here using the summary statistics method are, to 2 decimal places, identical to those computed using individual-level BCAC data and reported in Day et al. [2017]. As the outcome is binary, we do not provide an estimate of an indirect causal effect (see Discussion).

### Simulation study

To validate the utility of the multivariable Mendelian randomization method for estimating a direct causal effect, we performed a simulation analysis. We generated data on 10 genetic variants, a risk factor (*X*), mediator (*M*), and outcome (*Y*) for 10 000 individuals in a one-sample Mendelian randomization context. Full details of the simulation setup are provided in Supplementary Material A.3. Briefly, we considered eight different sets of values of the parameters *θ*_1_ (the causal effect of *X* on *M*), *θ*_2_ (the direct effect of *X* on *Y*), and *θ*_3_ (the effect of *M* on *Y*) – see Figure 2. The indirect effect of *X* on *Y* via *M* is *θ*_1_*θ*_3_, and the total effect of *X* on *Y* is *θ*_2_ +*θ*_1_*θ*_3_. We included scenarios where there is no direct effect, no indirect effect, a direct effect and a directionally concordant indirect effect, and a direct effect and a directionally discordant indirect effect. Parameters were chosen to take realistic values and cover a range of scenarios. 10 000 simulated datasets were generated for each choice of parameter values. Heterogeneity to ensure identification of the model was generated by additionally allowing the genetic variants to affect the mediator directly; these effects were drawn from a normal distribution with mean zero. Although this formally leads to pleiotropy and violation of the instrumental variable assumptions, it has been shown that such ‘balanced pleiotropy’ does not lead to bias in causal estimates [Bowden et al., 2015].

For each simulated dataset, we performed univariable Mendelian randomization analyses to estimate the total causal effect of the risk factor on the outcome, and multivariable Mendelian randomization for the direct causal effect not via the mediator. Each analysis was performed by weighted regression using the summarized data only (genetic associations with the risk factor, mediator, and outcome: beta-coefficients plus standard errors). We assumed that all genetic variants were uncorrelated (no linkage disequilibrium); their distributions in the data-generating model were independent. This assumption can be relaxed using generalized weighted linear regression as described elsewhere [Burgess et al., 2016].

Table 1 shows mean estimates of the total and direct effects, mean bias and standard deviations of the estimates, and coverage of the 95% confidence interval (the proportion of confidence intervals that include the true value of the parameter). The standard errors for the causal estimates were adjusted for underdispersion (residual standard error in the regression model less than 1) as described in the software code. No correction for overdispersion was applied [Burgess and Thompson, 2017]. The Monte Carlo error (uncertainty due to the limited number of simulations) was around 0.001 for each mean estimate, and 0.2% for the coverage proportion. We see that mean univariable Mendelian randomization estimates are similar to the total causal effect, whereas mean multivariable Mendelian randomization estimates are similar to the direct causal effect in each scenario considered. Bias in the mean estimates is small throughout, and is likely to be due to weak instrument bias arising from the limited strength of the genetic variants [Burgess et al., 2011] (no bias was observed on repeating the simulation study with a sample size of 1 000 000 for a small number of simulated datasets). Bias was consistent in direction for the total effect, but varied in direction for the direct effect. Coverage rates were close to nominal levels (95%) throughout, except for when there was substantial weak instrument bias in estimates of the direct effect. There was no noticeable undercoverage resulting from the regression models failing to account for uncertainty in the genetic associations with the risk factor or mediator. Further results in Supplementary Material A.4 indicate that these findings hold even when there are bidirectional effects of the risk factor on the mediator and vice versa (as may be the case for age at menarche and BMI).

**Table 1:** Mean, bias, standard deviation (SD), and coverage of 95% confidence interval (%) of nivariable and multivariable Mendelian randomization estimates across 10 000 simulated datasets r different mediation scenarios (*X* = risk factor, *M* = mediator, *Y* = outcome).

| $\theta_1$<br>($X \rightarrow M$) | $\theta_2$<br>($X \rightarrow Y$) | $\theta_3$<br>($M \rightarrow Y$) | Total effect<br>($\theta_2 + \theta_1\theta_3$) | Direct effect<br>($\theta_2$) | Univariable (total effect) | | | | Multivariable (direct effect) | | | |
| --- | --- | --- | --- | --- | --- | --- | --- | --- | --- | --- | --- | --- |
|  |  |  |  |  | Mean | Bias | SD | Coverage | Mean | Bias | SD | Coverage |
| 0.3 | 0.2 | 1 | 0.5 | 0.2 | 0.518 | 0.018 | 0.166 | 94.5 | 0.194 | -0.006 | 0.059 | 94.4 |
| 0.3 | 0.2 | -1 | -0.1 | 0.2 | -0.098 | 0.012 | 0.154 | 94.6 | 0.195 | -0.005 | 0.058 | 94.6 |
| 0.3 | -0.2 | 1 | 0.1 | -0.2 | 0.114 | 0.014 | 0.165 | 94.7 | -0.206 | -0.006 | 0.057 | 95.0 |
| -0.3 | -0.2 | 1 | -0.5 | -0.2 | -0.480 | 0.020 | 0.167 | 94.8 | -0.179 | 0.021 | 0.057 | 93.2 |
| 0.0 | 0.2 | 1 | 0.2 | 0.2 | 0.217 | 0.017 | 0.167 | 94.6 | 0.208 | 0.008 | 0.047 | 94.2 |
| 0.3 | 0.2 | 0 | 0.2 | 0.2 | 0.208 | 0.008 | 0.045 | 94.4 | 0.195 | -0.005 | 0.057 | 97.3 |
| 0.3 | 0.0 | 1 | 0.3 | 0.0 | 0.318 | 0.018 | 0.167 | 94.6 | -0.005 | -0.005 | 0.058 | 95.1 |
| -0.2 | 0.2 | 1 | 0.0 | 0.2 | 0.015 | 0.015 | 0.166 | 94.8 | 0.216 | 0.016 | 0.051 | 93.7 |

[Table 1 should appear around here]

## Discussion

In this paper, we have demonstrated how summarized data on genetic associations can be used to investigate causal mechanisms, in particular whether the causal effect of a complex risk factor on an outcome acts via a given mediator. Although the assumptions required for a genetic variant to be an instrumental variable are very stringent, in other ways, the requirements necessary to perform this analysis are quite flexible – only summarized data on genetic associations are required. This allows for the leverage of data from large-scale GWAS consortia. As with two-sample Mendelian randomization [Pierce and Burgess, 2013], the summarized data methods described here do not require the genetic associations with the risk factor, mediator and outcome to be measured in the same individuals. For example, Eppinga et al. used genetic variants to investigate the effect of resting heart rate on mortality in UK Biobank [Eppinga et al., 2016]. As a sensitivity analysis, they adjusted the genetic associations with the outcome for some covariates using individual-level data to assess whether the effect of resting heart rate was mediated via any of those variables. Additionally, they adjusted for genetic associations with lipid fractions using the multivariable Mendelian randomization approach outlined here, as lipid measurements are currently not available in the dataset. Combining summary statistics from different sources is also important in the example of age at menarche and breast cancer here, as BMI measurements for breast cancer cases were only available post-diagnosis. These measurements would likely be influenced by the disease process, as well as by treatment and lifestyle changes. It is therefore preferable here to estimate the effects of the genetic variants on BMI in a separate dataset.

### Compatibility of datasets

When using genetic associations from multiple datasets in a two-sample Mendelian randomization setting, ideally the associations should be estimated on samples from the same underlying population. This is particularly important with regard to ethnicity, as different linkage disequilibrium structures can mean that genetic variants may be associated with the risk factor in one population and not in another, or be valid instruments in one population but not in another. Ideally, genetic associations should not be adjusted for covariates apart from principal components of ancestry, particularly if these covariates may be on causal pathways relating to the risk factor, mediator or outcome. It is also important to ensure that genetic associations with the risk factor and mediator are estimated in individuals who have not had disease events, so that these associations are not influenced by reverse causation. However, even if associations are estimated in different datasets (say, associations with the risk factor are measured in 20-year olds and associations with the mediator in 50-year olds, or vice versa), as genetic variants typically influence variables across the whole lifecourse, inferences from Mendelian randomization for the causal null hypothesis should still be qualitatively valid, even if the parametric assumptions necessary for causal estimation are not satisfied [Burgess et al., 2016]. In any case, as Mendelian randomization estimates represent the effect of changing people’s genetic variants at conception, causal estimates from Mendelian randomization should not be interpreted too literally as the expected impact of intervening on the risk factor in practice [Burgess et al., 2012]. These issues are discussed in greater detail in Burgess et al. [2016] and Bowden et al. [2017].

In the context of mediation, potential inconsistencies in genetic association estimates from different sources are more important. In univariable Mendelian randomization, if the genetic associations with the risk factor are misspecified, then the inverse-variance weighted estimate is still a weighted sum of the genetic associations with the outcome, and should differ from zero when the instrumental variable assumptions are satisfied if and only if there is a causal effect of the risk factor on the outcome. However in multivariable Mendelian randomization, if genetic associations with the mediator are misspecified, then adjustment for genetic associations with the mediator may not fully attenuate the coefficient in the weighted regression for the effect of the risk factor even in the case of complete mediation. Multiplying genetic associations by a constant would not affect the significance of coefficients in the weighted regression, hence any differences between populations that would lead to consistent over-or underestimation of genetic associations for all variants should not influence inferences from the methods presented here. However, differences that lead to inconsistent over-or underestimation of genetic associations would adversely affect causal inferences. Therefore, genetic associations should be estimated in as similar populations as possible.

### Binary variables and non-linear relationships

It is common for the outcome in a Mendelian randomization investigation to be a binary variable, such as disease status. In this case, typically genetic associations are obtained from logistic regression, and represent log odds ratios. Odds ratios are non-collapsible, meaning that they do not average intuitively, and they depend on the choice of covariate adjustment even in the absence of confounding (so conditional odds ratios differ in magnitude to marginal odds ratios) [Greenland et al., 1999]. This means that differences between causal estimates from equations (1) and (2) may arise due to non-collapsibility rather than mediation. However, these differences are likely to be slight [Burgess, 2017]. In practice, as in the applied example considered in this paper, we would recommend providing estimates of the total and direct effects, but not the indirect effect, as calculation of the indirect effect relies on the linearity of the relationships that cannot occur with a binary outcome. The total and direct effects still have interpretations as population-averaged causal effects (conditional on the mediator for the direct effect), representing the average change in the outcome resulting from intervening on the population distribution of the risk factor (while keeping the mediator constant for the direct effect) [Burgess and CHD CRP Genetics Collaboration, 2013]. Substantial differences between these estimates would still be informative about the causal pathway from the risk factor to the outcome.

Similarly, if there is a non-linear relationship between the risk factor and outcome, the causal effects still have an interpretation as population-averaged causal effects, representing the average change in the outcome resulting from intervening on the population distribution of the risk factor [Burgess et al., 2014]. Again, we would recommend reporting a total effect and a direct effect, but not an indirect effect.

In conclusion, we hope that the methods outlined in this manuscript will be used widely in assessing and understanding causal pathways and mechanisms.

## Acknowledgements

This study would not have been possible without the contributions of the following: Per Hall (COGS); Douglas F. Easton, Paul Pharoah, Kyriaki Michailidou, Manjeet K. Bolla, Qin Wang (BCAC), Andrew Berchuck (OCAC), Rosalind A. Eeles, Douglas F. Easton, Ali Amin Al Olama, Zsofia Kote-Jarai, Sara Benlloch (PRACTICAL), Georgia Chenevix-Trench, Antonis Antoniou, Lesley McGuffog, Fergus Couch and Ken Offit (CIMBA), Joe Dennis, Alison M. Dunning, Andrew Lee, and Ed Dicks, Craig Luccarini and the staff of the Centre for Genetic Epidemiology Laboratory, Javier Benitez, Anna Gonzalez-Neira and the staff of the CNIO genotyping unit, Jacques Simard and Daniel C. Tessier, Francois Bacot, Daniel Vincent, Sylvie LaBoissi`ere and Frederic Robidoux and the staff of the McGill University and Génome Québec Innovation Centre, Stig E. Bojesen, Sune F. Nielsen, Borge G. Nordestgaard, and the staff of the Copenhagen DNA laboratory, and Julie M. Cunningham, Sharon A. Windebank, Christopher A. Hilker, Jeffrey Meyer and the staff of Mayo Clinic Genotyping Core Facility

Funding for the iCOGS infrastructure came from: the European Community’s Seventh Framework Programme under grant agreement number 223175 (HEALTH-F2-2009-223175) (COGS), Cancer Research UK (C1287/A10118, C1287/A 10710, C12292/A11174, C1281/A12014, C5047/A8384, C5047/A15007, C5047/A10692), the National Institutes of Health (CA128978) and Post-Cancer GWAS initiative (1U19 CA148537, 1U19 CA148065 and 1U19 CA148112 - the GAME-ON initiative), the Department of Defence (W81XWH-10-1-0341), the Canadian Institutes of Health Research (CIHR) for the CIHR Team in Familial Risks of Breast Cancer, Komen Foundation for the Cure, the Breast Cancer Research Foundation, and the Ovarian Cancer Research Fund.

Stephen Burgess is supported by Sir Henry Dale Fellowship jointly funded by the Wellcome Trust and the Royal Society (Grant Number 204623/Z/16/Z).

## Supplementary Material

### A.1 Technical discussion about estimation of indirect and direct effects

There are several versions of direct and indirect effects. We present definitions using counter-factual terminology, using potential values of the outcome *Y* (*x, m*), representing the outcome which would be observed if *X* were set (by intervention) to *x* and *M* were set to *m*, and potential values of the mediator *M* (*x*), the value taken by the mediator if *X* were set to *x*. All effects are given on the difference scale; with a binary outcome, effects on a relative risk or odds ratio scale can also be defined, but the decomposition is more complex [VanderWeele and Vansteelandt, 2010; Kaufman, 2010]. This text is adapted from Burgess et al. [2015].

A total effect is defined as the effect of a change in the exposure from, say, *X* = *x* to *X* = *x* + 1. It comprises the effects of the change in the exposure, and the change in the mediator as a result of the change in the exposure:

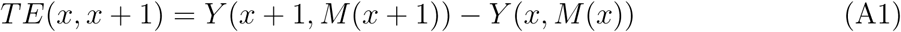

A controlled direct effect is defined as the effect of a change in the exposure keeping the mediator fixed at a given level, say *M* = *m* [Robins and Greenland, 1992; Pearl, 2001]. The controlled direct effect may depend on the choice of *m*:

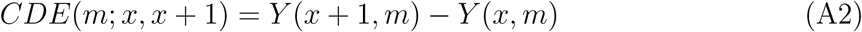

A natural direct effect is defined as the effect of a change in the exposure with the mediator fixed at the level it would naturally take if the exposure were fixed at a given level, say *X* = *x*:

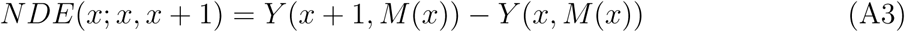

A natural indirect effect is defined as the effect of a change in the mediator from the value it would naturally take if the exposure were unchanged to the level it would take if the exposure were changed. The exposure itself is kept fixed at a given level, say *X* = *x* + 1:

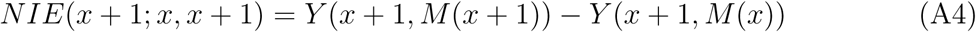

In the linear case, the natural direct and indirect effects represent a decomposition of the total effect, in that *TE*(*x, x* + 1) = *NDE*(*x*; *x, x* + 1) + *NIE*(*x* + 1; *x, x* + 1) (or alternatively *TE*(*x, x* + 1) = *NDE*(*x* + 1; *x, x* + 1) + *NIE*(*x*; *x, x* + 1)). Under the condition:

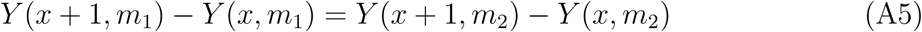

for all values of *M* = *m*_1_*, m*_2_, and for all individuals, the controlled direct effect is equal to the natural direct effect [Robins and Greenland, 1992]. The natural direct effect has a clearer intuitive interpretation as a measure of mediation than the controlled direct effect. However, it is not possible to conceive of an experiment which would produce the natural direct effect, as the quantity requires the outcome if the exposure were set at two different levels (for example, in *NDE*(*x*; *x, x* + 1), *Y* (*x* + 1*, M* (*x*)) requires *X* = *x* + 1 for *Y*, but *X* = *x* for *M*). This is known as a “cross-world” quantity, as setting the exposure to two different values is only possible in two different worlds [Richardson and Robins, 2013].

As we argue in Burgess et al. [2015], we would regard the controlled direct effect as the quantity that is targeted by mediation analysis with instrumental variables, as this is what would be obtained if we were to intervene separately on the risk factor and mediator. As we assume that all relationships between variables are linear and there is no effect heterogeneity, the natural and controlled direct effects are equal, and hence we refer to a ‘direct effect’ throughout this manuscript without further qualification.

### A.2 Software code

We provide R code to implement the methods discussed in this paper. The associations of the genetic variants with the risk factor are denoted betaXG with standard errors sebetaXG. The associations of the genetic variants with the mediator are denoted betaMG with standard errors sebetaMG. The associations of the genetic variants with the outcome are denoted betaYG with standard errors sebetaYG. When variables are continuous, these associations are typically estimated using linear regression.

Estimation of the total causal effect using summarized data:

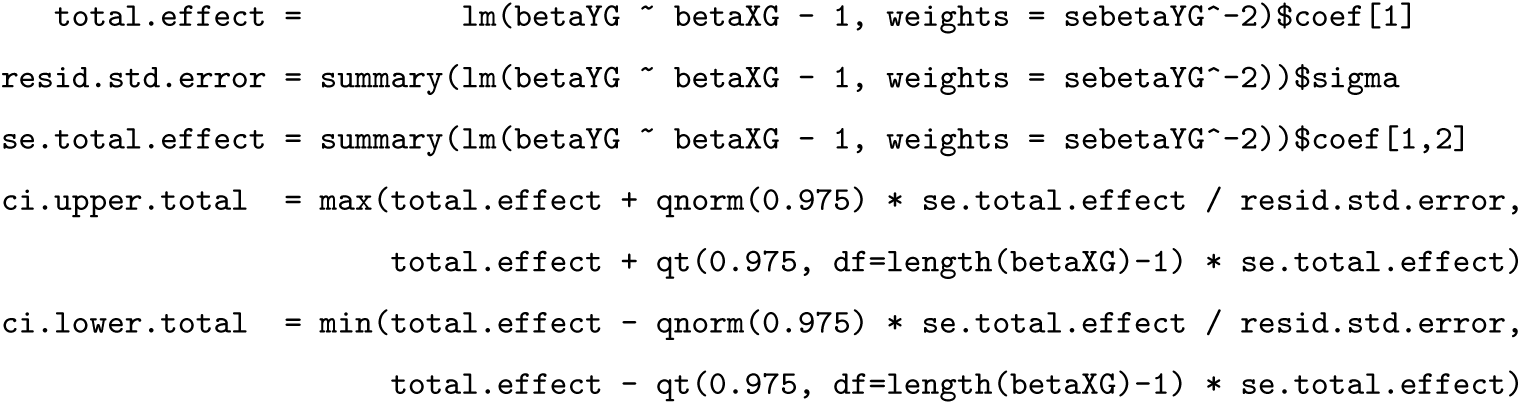

The weighted regression model for estimating the total effect is equivalent to a meta-analysis of the variant-specific causal estimates. Setting the residual standard error as 1 is equivalent to a fixed-effect assumption in the meta-analysis formula [Thompson and Sharp, 1999]. If there is no heterogeneity between the causal estimates identified by the individual variants, then the residual standard error should tend to 1 asymptotically. If the estimate of the residual standard error is greater than 1 (overdispersion), then we do not correct for this; this is equivalent to a (multiplicative) random-effects meta-analysis [Burgess and Thompson, 2017]. This would occur if different genetic variants identify different causal estimates (say, different variants influence the risk factor via different mechanisms). However, there is no biological rationale for underdispersion (residual standard error estimate is less than 1). Hence, we correct for underdispersion by dividing the standard error for the total effect by the residual standard error.

The multiplicative random-effects analysis fits the following model, with *ϕ* representing the residual standard error:

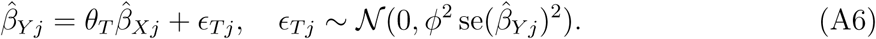

For a fixed-effect analysis, the residual standard error is assumed to be known; hence it is appropriate to use a normal distribution for inferences. For a random-effect analysis, as the residual standard error (the overdispersion parameter *ϕ*) is estimated rather than known, a t-distribution should be used for making inferences. In the confidence intervals, we take the upper bound to be the maximum of the bounds based on the fixed-effect and random-effect analyses; similarly for the lower bound as the minimum. This ensures that confidence intervals are no wider than they would be from a fixed-effect analysis, but that under-precision is not doubly penalized (by setting the residual standard error to be 1, and then using a t-distribution for inferences).

Estimation of the direct causal effect using summarized data:

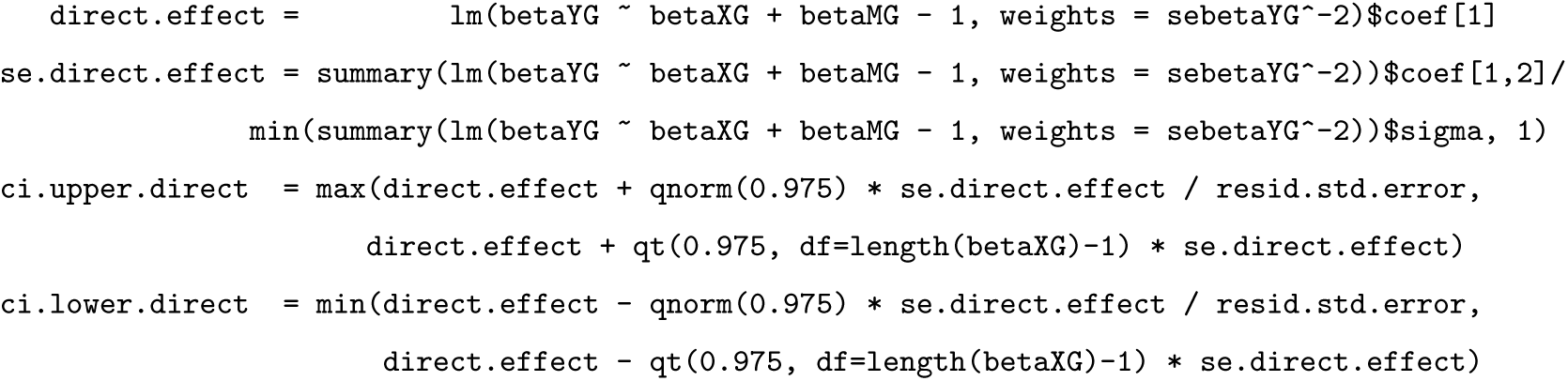

As the additional term in the regression analysis for the estimate of the direct effect lowers the residual standard error, we take the estimated residual standard error from the regression model for the total causal effect. This is because we want this term to represent overdispersion in the genetic associations with the outcome, not the residual associations after adjustment. Hence the t-distribution for making inferences is still on *J -* 1 degrees of freedom.

If the outcome is binary, then genetic associations with the outcome are typically estimated using logistic regression. Beta-coefficients from logistic regression can be used in the estimation of direct and indirect effects, but the precise magnitude of effect estimates should not be over-interpreted, as odds ratios suffer from non-collapsibility when the rare disease assumption is not applicable (instrumental variable estimates represent population-averaged causal effects, which are not the same as subject-specific causal effects on the odds ratio scale, hence the indirect and direct effects may not precisely sum to give the total effect). Therefore in the applied example in this paper, we do not report an indirect effect.

With correlated variants, this correlation can be accounted for by generalized weighted linear regression [Burgess et al., 2016]. We assume that rho is the matrix of correlations between genetic variants:

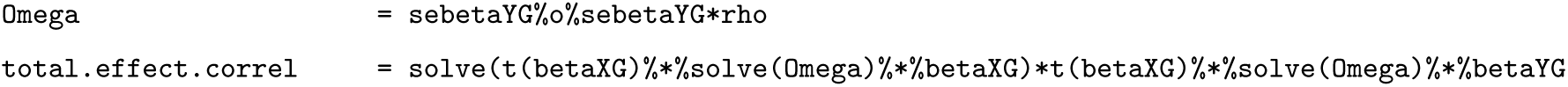
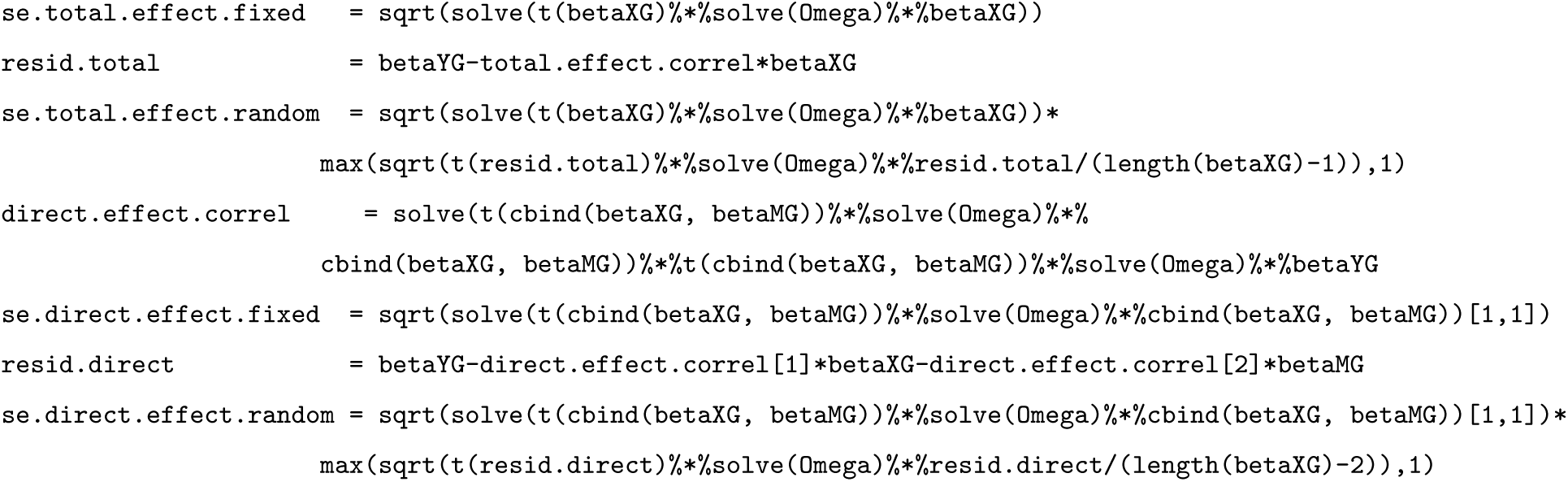

Standard errors are given corresponding to both fixed-effect and random-effects assumptions.

### A.3 Additional details of simulation study

For the simulation study in the paper, the risk factor *X* was generated as:

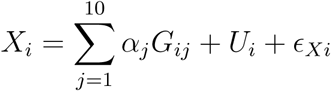

where *G*_*ij*_ is the number of variant alleles for genetic variant *j*, *U* is a confounder, *ϵ*_*Xi*_ is an independent error term. The number of variant alleles for each variant was drawn from a binomial distribution with 2 trials and probability 0.3, representing a single nucleotide polymorphism with minor allele frequency 0.3. The genetic effects on the risk factor *α*_*j*_ were generated from a normal distribution with mean 0.2 and variance 0.12. The variants in total explained on average 5.1% of the variance in the risk factor, corresponding to an average F statistic of 53.5 with a sample size of 10 000. The confounder *U* and all error terms (*ϵ*_*X*_*, ϵ*_*M*_ *, ϵ*_*Y*_) were drawn from independent standard normal distributions. The mediator *M* was generated as:

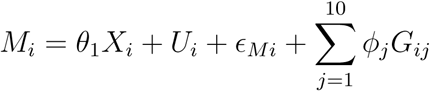

where *θ*_1_ is the causal effect of *X* on *M*, and *ϕ*_*j*_ are direct effects of the genetic variants on the mediator. These effects are included in the simulation model to ensure that the direct effect is identified, as otherwise genetic associations with the risk factor and mediator would be perfectly correlated for large sample sizes, leading to unstable estimates of the direct effect. The *ϕ*_*j*_ parameters were generated from a normal distribution with mean zero and variance 0.12. The outcome *Y* was generated as:

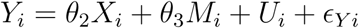

where *θ*_2_ is the direct effect of *X* on *Y*, and *θ*_3_ is the effect of *M* on *Y*. The indirect effect of *X* on *Y* via *M* is *θ*_1_*θ*_3_, and the total effect of *X* on *Y* is *θ*_2_ + *θ*_1_*θ*_3_. In total, 10 000 simulated datasets were generated for each choice of parameter values.

We experimented with different values of the variance of the *ϕ*_*j*_ parameters in the datagenerating model. Results are shown in Supplementary Table A1. When there was low heterogeneity, estimates were more variable and bias from weak instruments was more pronounced. This is expected, as the associations with the risk factor and mediator are increas-ingly collinear as the heterogeneity decreases. To demonstrate that the bias is an artifact of limited sample size (so called ‘weak instrument bias’), we repeated the simulation with 100 000 participants (100 iterations per scenario only). As expected, bias did not decrease when there was no heterogeneity, as the collinearity problem does not disappear with increasing sample sizes in this case. However, in all other cases, increasing the sample size decreased bias sharply.

**Table A1:** Mean (standard deviation) of multivariable Mendelian randomization estimates of the direct effect *θ*_2_ across 10 000 simulated datasets (100 datasets for larger sample size) for different values of the variance of the heterogeneity parameters *ϕ*.

| Sample size: 10 000 |  |  |  |  |  |  |
| --- | --- | --- | --- | --- | --- | --- |
| $\theta_1$ | $\theta_2$ | $\theta_3$ | $\text{var}(\phi) = 0$ | $\text{var}(\phi) = 0.05^2$ | $\text{var}(\phi) = 0.1^2$ | $\text{var}(\phi) = 0.2^2$ |
| 0.3 | 0.2 | 1 | 0.054 (0.110) | 0.165 (0.077) | 0.196 (0.055) | 0.203 (0.050) |
| 0.3 | 0.2 | -1 | 0.052 (0.115) | 0.165 (0.076) | 0.195 (0.056) | 0.204 (0.050) |
| 0.3 | -0.2 | 1 | -0.343 (0.113) | -0.235 (0.071) | -0.205 (0.059) | -0.194 (0.050) |
| -0.3 | -0.2 | 1 | -0.049 (0.101) | -0.153 (0.073) | -0.181 (0.058) | -0.187 (0.052) |
| 0.0 | 0.2 | 1 | 0.205 (0.041) | 0.207 (0.048) | 0.208 (0.048) | 0.207 (0.048) |
| 0.3 | 0.2 | 0 | 0.053 (0.106) | 0.168 (0.074) | 0.196 (0.058) | 0.203 (0.053) |
| 0.3 | 0.0 | 1 | -0.146 (0.113) | -0.035 (0.071) | -0.004 (0.059) | 0.003 (0.050) |
| -0.2 | 0.2 | 1 | 0.302 (0.076) | 0.235 (0.057) | 0.213 (0.050) | 0.210 (0.048) |
| Sample size: 100 000 |  |  |  |  |  |  |
| $\theta_1$ | $\theta_2$ | $\theta_3$ | $\text{var}(\phi) = 0$ | $\text{var}(\phi) = 0.05^2$ | $\text{var}(\phi) = 0.1^2$ | $\text{var}(\phi) = 0.2^2$ |
| 0.3 | 0.2 | 1 | 0.053 (0.092) | 0.191 (0.027) | 0.200 (0.019) | 0.201 (0.016) |
| 0.3 | 0.2 | -1 | 0.051 (0.114) | 0.194 (0.030) | 0.195 (0.019) | 0.202 (0.016) |
| 0.3 | -0.2 | 1 | -0.341 (0.098) | -0.206 (0.028) | -0.197 (0.016) | -0.198 (0.015) |
| -0.3 | -0.2 | 1 | -0.049 (0.087) | -0.191 (0.027) | -0.197 (0.020) | -0.202 (0.017) |
| 0.0 | 0.2 | 1 | 0.199 (0.012) | 0.202 (0.016) | 0.199 (0.016) | 0.200 (0.016) |
| 0.3 | 0.2 | 0 | 0.055 (0.106) | 0.196 (0.027) | 0.199 (0.019) | 0.200 (0.017) |
| 0.3 | 0.0 | 1 | -0.136 (0.099) | -0.005 (0.027) | 0.003 (0.018) | 0.000 (0.017) |
| -0.2 | 0.2 | 1 | 0.296 (0.072) | 0.206 (0.018) | 0.200 (0.018) | 0.200 (0.016) |

### A.4 Additional simulation scenario: bidirectional causal effects between risk factor and mediator

In the applied example, it may be that as well as the risk factor having a causal effect on the mediator, that the mediator also has a causal effect on the risk factor. To consider this scenario, we simulate causal effects in both directions and consider Mendelian randomization and multivariable Mendelian randomization estimates. The data-generating model is:

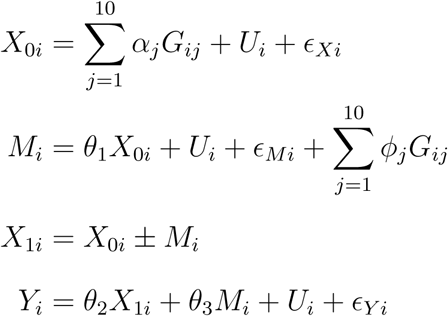

This is the same as the previous data-generating model, except that we first generate *X*_0*i*_ and then generate a second risk factor variable *X*_1*i*_ that has a causal effect from the mediator. These could be thought of as values of the risk factor at different time points. We consider cases where the mediator has a positive and a negative effect on the risk factor. All other aspects of this simulation are the same as the original.

Results are shown in Supplementary Table A2. The total effect varies depending on whether the effect of the mediator on the risk factor is positive or negative, and is not simply an estimate of *θ*_2_ + *θ*_1_*θ*_3_ (as there are additional components of the total effect via the effect of the mediator on the risk factor). However, the direct effect as estimated by multivariable Mendelian randomization is invariant to any bidirectional effect. Therefore the direct effect of age at menarche on breast cancer risk not via BMI can be estimated using multivariable Mendelian randomization whether or not there is a bidirectional relationship between age at menarche and BMI.

**Table A2:** Mean of univariable and multivariable Mendelian randomization estimates across 10 000 simulated datasets for different mediation scenarios with positive and negative bidirectional effect of the mediator on the risk factor.

| $\theta_1$ | $\theta_2$ | $\theta_3$ | Positive effect | | Negative effect | |
| --- | --- | --- | --- | --- | --- | --- |
|  |  |  | Univariable | Multivariable | Univariable | Multivariable |
| 0.3 | 0.2 | 1 | 0.525 | 0.195 | 0.222 | 0.195 |
| 0.3 | 0.2 | -1 | -0.103 | 0.194 | 0.173 | 0.194 |
| 0.3 | -0.2 | 1 | 0.123 | -0.204 | -0.169 | -0.204 |
| -0.3 | -0.2 | 1 | -0.195 | -0.180 | -0.504 | -0.180 |
| 0.0 | 0.2 | 1 | 0.381 | 0.208 | 0.045 | 0.208 |
| 0.3 | 0.2 | 0 | 0.209 | 0.195 | 0.197 | 0.195 |
| 0.3 | 0.0 | 1 | 0.323 | -0.005 | 0.017 | -0.005 |
| -0.2 | 0.2 | 1 | 0.273 | 0.217 | -0.060 | 0.217 |

## Figure titles and legends

Figure 1. Title: Graphical representation of mediation scenario. Caption: Total effect of risk factor on outcome comprises an indirect effect (hollow arrows) via mediator, and a direct effect (solid arrow) via other pathways.

Figure 2. Title: Graphical representation of relationships between variables. Caption: Graphical diagram of relationships between risk factor (*X*), mediator (*M*), outcome (*Y*) and genetic variant (*G*_*j*_). Causal relationships between variables are indicated by solid lines. Associations of the genetic variant are indicated by dashed lines. The direct effect *θ*_*D*_ = *θ*_2_. The indirect effect *θ*_*I*_ = *θ*_1_*θ*_3_. The total effect *θ*_*T*_ = *θ*_*D*_ + *θ*_*I*_ = *θ*_2_ + *θ*_1_*θ*_3_.

**Table title and legend**

Table 1. Title: Simulation study results. Caption: Mean, bias, standard deviation (SD), and coverage of 95% confidence interval (%) of univariable and multivariable Mendelian randomization estimates across 10 000 simulated datasets for different mediation scenarios (*X* = risk factor, *M* = mediator, *Y* = outcome).

